# GammaGateR: semi-automated marker gating for single-cell multiplexed imaging

**DOI:** 10.1101/2023.09.20.558645

**Authors:** Jiangmei Xiong, Harsimran Kaur, Cody N Heiser, Eliot T McKinley, Joseph T Roland, Robert J Coffey, Martha J Shrubsole, Julia Wrobel, Siyuan Ma, Ken S Lau, Simon Vandekar

## Abstract

**Motivation:** Multiplexed immunofluorescence (mIF) is an emerging assay for multichannel protein imaging that can decipher cell-level spatial features in tissues. However, existing automated cell phenotyping methods, such as clustering, face challenges in achieving consistency across experiments and often require subjective evaluation. As a result, mIF analyses often revert to marker gating based on manual thresholding of raw imaging data.

**Results:** To address the need for an evaluable semi-automated algorithm, we developed GammaGateR, an R package for interactive marker gating designed specifically for segmented cell-level data from mIF images. Based on a novel closed-form gamma mixture model, GammaGateR provides estimates of marker-positive cell proportions and soft clustering of marker-positive cells. The model incorporates user-specified constraints that provide a consistent but slide-specific model fit. We compared GammaGateR against the newest unsupervised approach for annotating mIF data, employing two colon datasets and one ovarian cancer dataset for the evaluation. We showed that GammaGateR produces highly similar results to a silver standard established through manual annotation. Furthermore, we demonstrated its effectiveness in identifying biological signals, achieved by mapping known spatial interactions between CD68 and MUC5AC cells in the colon and by accurately predicting survival in ovarian cancer patients using the phenotype probabilities as input for machine learning methods. GammaGateR is a highly efficient tool that can improve the replicability of marker gating results, while reducing the time of manual segmentation.

**Availability and Implementation:** The R package is available at https://github.com/JiangmeiRubyXiong/GammaGateR.

**Contact:** Please address correspondence to.

**Key Points:**

- GammaGateR is the first semi-automated marker gating tool for mIF image, and it will help to diminish the inconsistency with manual marker gating.
- With novel cfGMM, GammaGateR can fit flexibly across slides with different distributions and incorporate biology priors.
- GammaGateR is proven to reveal credible prognostic information, and it can quantify known findings in tumor cell populations.

## 1. Introduction

Multiplexed immunofluorescence (mIF) imaging is a recently developed spatial proteomic assay that allows the investigation of a tissue microenvironment for many markers and high spatial resolution [1–4]. mIF images are obtained through cyclic imaging with up to 60 marker channels, each identifying a specific protein. mIF is advantageous over other single-cell assays that disaggregate the tissue because it can reveal insights about the spatial interactions between tissue and cell types *in situ* and offers a subcellular spatial resolution that is higher than other spatial assays. mIF has already revealed valuable spatial insights into the tumor microenvironment [5–8]. For example, recent findings demonstrate differences in immune cell infiltration of colorectal precancerous polyps such as a higher concentration of CD8+ T-cells in the epithelium of sessile serrated lesions (SSL) than that of conventional adenomas and higher CD68+ macrophages concentration at the luminal surface in SSL, but more randomly distributed in adenomas [5].

Before important spatial insights can be gleaned using statistical methods [9–12], mIF images undergo an intensive preprocessing pipeline to obtain single-cell measurements. While there are various steps included in the pipeline such as image registration, single-cell segmentation, quantification, and batch correction [13–16], cell phenotyping is typically the final step before downstream analyses on the cell-level data, similarly to other single-cell assays. Cell phenotyping identifies individual cell phenotypes from the measured marker expression values of the cell and directly affects the subsequent cell population analysis results.

The two most common approaches for cell phenotyping in mIF are manual gating and graph-based multivariate clustering. In manual gating, each sample is visualized separately to determine a threshold, and super-threshold cells are labeled as marker positive. This procedure is repeated for all marker channels and slides, and the phenotypes are determined by combining combinations of marker-positive cells [5,8,17]. Alternatively, multivariate graph-based clustering is adapted from other single-cell assays [18,19]. This approach first performs cell clustering, then assigns a phenotype to each cell group based on their average expression profile. Multivariate graph-based clustering is implemented with various modifications across many software packages [20–22]. Unfortunately, both methods are labor intensive, and their accuracy suffers from image noise and spatial artifacts in mIF images that cause marker expression histograms to appear continuous or uni-modal . As a result, both phenotyping methods possess shortcomings that cannot be ignored. On one hand, manual gating can be subjective. On the other hand, graph-based clustering results are prone to over-clustering and producing poor separation between clusters [23,24].

The challenges described above are well recognized and there are a few methods and software developed that attempt to automate cell phenotyping for mIF images [22,25–28]. For example, CellSighter is a recently proposed supervised deep-learning algorithm for cell phenotyping that requires a “gold standard” training dataset [29]. Another recent solution, ASTIR (Automated assignment of cell identity from single-cell multiplexed imaging and proteomic data), is a fast unsupervised approach that defines cell phenotypes from segmented cell-level data by using a neural network-based mixture model assuming a multivariate log-normal distribution [30].

Instead of binary outputs like in classification methods, ASTIR returns posterior probabilities of different cell types for each cell. This type of output is advantageous because it offers more information than nominal cell types and leaves cell labeling to the clinician’s discretion. Lastly, Ahmadian et al. treat the analysis as a pixel classification problem and design a single-step framework for mIF phenotyping that is integrated with other preprocessing steps [31].

Nevertheless, inconsistencies persist in the results rendered by these learning-based methods when applied across markers, slides, batches, and datasets. These inconsistencies result from the immense variation in the cell-level distribution of phenotyping markers that are often too nuanced to be removed by existing batch correction methods [13–15]. For these reasons, it is difficult to fully automate the cell phenotyping process, despite the availability of automated tools, and manual gating is still used to perform cell phenotyping because it is easy to visualize and evaluate the quality of the phenotype [5,8].

Because automated methods cannot be run without evaluation and supervised methods require a gold-standard dataset, no method is truly fully automated. As a solution, we develop an explicitly *semi-automated* algorithm called GammaGateR. GammaGateR allows the user to easily perform cell phenotyping, visualize results, and conduct interpretable quality control while reducing manual labor. Based on a novel closed-form Gamma mixture model (cfGMM), GammaGateR is a probabilistic model that is fitted to each channel and slide separately, and outputs positive-component probabilities for each marker. These can then be easily thresholded and combined for semi-automated marker gating or input directly into downstream analysis.

GammaGateR has important technical advantages, including 1) improved computation time and model convergence due to its novel closed-form property, and 2) high consistency and reproducibility for phenotyping results across mIF data batches due to incorporation of parameter boundary constraints. In applications on real-world mIF data, we find that GammaGateR has fast and consistent results across many slides and markers. We provide an open-source implementation of our method in the new GammaGateR R package (https://github.com/jiangmeirubyxiong/gammagater).

In this paper, we describe the cfGMM model, and evaluate its computational performance and statistical properties. We then compare the accuracy of the GammaGateR with the current state-of-the-art unsupervised cell-level phenotyping tool, ASTIR, in three datasets. Finally, we use GammaGateR outputs to compare spatial features of immune cell populations in tissue compartments between adenoma and serrated colon polyps.

## 2. Methods

### 2.1 Datasets and Preprocessing

We use three single-cell imaging datasets to evaluate model performance and demonstrate the use of the GammaGateR analysis pipeline (Table 1): the Colorectal Molecular Atlas Project (Colon MAP) dataset [5], the Spatial Colorectal Cancer (CRC) Atlas dataset [7] and Ovarian Cancer dataset [32,33]. Dataset-specific acquisition and processing are described below and in prior work [5,7,33]. After processing and prior to analysis, cell expression values were normalized by first mean division then log10 transformation to reduce slide-to-slide variation [14]. Data collection for the Colon MAP and CRC atlas was approved by the Institutional Review Board (IRB) at Vanderbilt University Medical Center and collection of the ovarian cancer dataset was approved by the IRB at the University of Colorado.

**Table 1:** Sample characteristics for each dataset.

| Dataset | Number of slides | Number of patients | Number of Cells | Number of Marker Channels | Manual Marker Label Included | Patient Survival Information Included | Tumor Mask |
| --- | --- | --- | --- | --- | --- | --- | --- |
| Colon Map | 42 | 42 | 2,244,117 | 33 | Yes | No | Yes |
| CRC Atlas | 16 | 16 | 9,309,130 | 33 | Yes | No | No |
| Ovarian Cancer | 2 | 128 | 1,610,431 | 9 | No | Yes | Yes |

#### Colon MAP DataCe

The Colon MAP data consists of precancerous samples including conventional adenomas and SSL. SSLs are more often found in the proximal colon, represent only 10-20% of all polyps, and exhibit higher cytotoxic immune cell infiltration [5]. The goal of this study was to characterize differences in the microenvironment of these two types of polyps. Data collection and processing are as described in [5]. Briefly, imaging was performed on a GE IN Cell Analyzer 2500 using the Cell DIVE platform and acquired at x200 magnification, with exposure times determined manually for each antibody. Details on antibodies, staining sequence, and exposure times are given in their previous report [5]. Single-cell segmentation for the colon datasets was performed using MIRIAM, a multichannel machine learning single-cell segmentation algorithm designed and evaluated for human colon and human breast carcinoma [16]. Single-cell channel quantification was performed by taking the median value within each cell region and these values were normalized and used for subsequent analyses [14]. Epithelial and stromal tissue compartments were estimated using Leiden clustering and the tumor region of each sample was manually identified.

#### CRC Atlas Data

The CRC atlas dataset includes 16 samples that represent various stages of human colon cancer [7]. The goal of this study was to investigate the co-evolution of tumor and immune microenvironments in microsatellite-stable and chromosomally-unstable tumors. A molecular profiling assay was performed using mIF with 33 marker channels. Single-cell segmentation, channel quantification, and normalization are performed as in the Colon MAP dataset.

#### Ovarian Cancer Data

The Ovarian Cancer data includes multiplexed immunohistochemistry (IHC) images from a tissue microarray of 128 patients with high-grade serous carcinoma [33,34]. The data were collected using Vectra Automated Quantitative Pathology Systems (Akoya Biosciences) at 20x resolution and stained with antibodies specific for CD8, CD68,cytokeratin, CD3, and CD19. Preprocessing including single-cell segmentation was performed using inForm software version 2.3 (Akoya Biosciences). The data are freely available through the VectraPolaris Bioconductor Package [32] .

### 2.2 GammaGateR

#### Overview

The GammaGateR algorithm is unique to existing methods for its focus on parsimoniously modeling cell-level marker expression densities. This approach yields tailored-to-slide model estimation in cell-level mIF data where marker expression distributions can vary substantially across slides. The algorithm uses a zero-inflated two-component GMM to model marker expression for each slide. The Gamma mixture model naturally identifies marker-positive and marker-negative cell distributions and returns the probability of belonging to the marker-positive cell distribution for each cell. The returned probabilities can either be used directly in subsequent analysis or combined and dichotomized to define cell phenotypes. GammaGateR incorporates user-specified constraints to provide consistent model fit across a large number of slides. The model evaluation methods included in GammaGateR allow the user to evaluate the constraints and quality check results. The power source of GammaGateR is the closed-form Gamma mixture model, which is a novel approach to phenotyping for mIF data that makes it more computationally efficient than traditional GMMs.

#### Closed-form Gamma mixture model estimation

For mIF data, we use the GMM to fit cell marker expression values as a weighted sum of different probability distributions that represent unique cell populations [35]. The Gamma distribution is an excellent model for marker values because the domain of the Gamma distribution is strictly positive and it has the flexibility to model the varying skewed densities seen in mIF marker values (Figure 1.5.a). However, GMMs are not scalable for mIF image data, because they rely on computationally inefficient numerical methods to obtain the maximum likelihood estimator (MLE). The slow convergence of the MLE for the GMM makes it prohibitive to apply across a large number of channels, slides, and cells. As a solution, we develop a closed-form GMM (cfGMM; https://github.com/jiangmeirubyxiong/cfgmm) estimation procedure based on a recently developed estimator for the Gamma distribution [36]. In addition, to improve computational efficiency, the cfGMM has the benefit of allowing prior constraints on model parameters. With the cfGMM in GammaGateR, we enable the flexibility to include a biologically meaningful range for the mode of each component in the Gamma mixture model. This way, users of GammaGateR can restrict estimation to biologically meaningful values.

For the GammaGateR model, we assume nonzero marker values belong to two components representing marker positive and negative cells separately. Because mIF data can include zero values due to autofluorescence adjustment, we use a zero-inflated two-component Gamma mixture model:

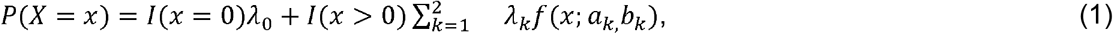

where *λ*_0_ represents the proportion of zeros, *λ*_l_ is the proportion of marker-negative cells, and *λ*_2_ is the proportion of marker-positive cells. *I*(·) denotes the indicator function and *f(x; a*_*k*,_*b*_*k*_) is the Gamma density function corresponding to the *k*th component, with *a*_*k*_ and *b*_*k*_ as its shape and scale parameters for the *k*th component and *m*_*k*_ *= (a*_*k*_ − 1)*b*_*k*_ as its mode. We assume that *m*_2_ > *m*_l_, that is, the component with higher mode corresponds to the marker-positive cells.Given values of *λ* _*k*_, *a*_*k*_ and *b*_*k*_, the posterior probability of a given cell being marker positive is

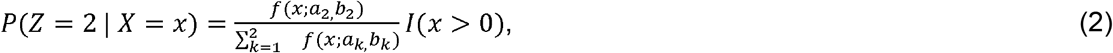

where *Z* is a random variable indicating the component membership of the given cell (0,1,2). In the extreme tails, the marker-positive probability might not always be higher than that of marker-negative, due to different variances of the two components’ Gamma distributions. Therefore, we apply a correction to the posterior probabilities to force them to be monotonic with respect to the marker values. Specifically, after the first crossing of the density curves of the two components, the density curve of the first component will be forced to be non-increasing, and the density curve of the second component will be forced to be non-decreasing. In addition to the posterior probability, GammaGateR also outputs the marginal probability of the observed marker value for the marker-positive component

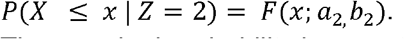

The marginal probability is monotonically increasing in the marker intensities and represents the probability that a marker positive cell is less than the given value, *x*.

We improve computation time of the classical GMM using a recently developed closed-form estimator for the Gamma distribution [36]. We replace the Gamma distribution, *f*, in equation (1) with the generalized gamma and derive the closed-form estimators for the GMM using the approach used for the Gamma distribution by Ye and Chen (2017),

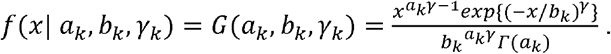

When *γ*_*k*_ = 1, Equation 1 is equal to the gamma distribution with shape *a*_*k*_ and scale *b*_*k*_. Closed-form estimators of *a*_*k*_ and *b*_*k*_ are obtained by differentiating Equation 1 with respect to *b*_*k*_ and *γ*_*k*_, setting the derivative to 0, solving for *a*_*k*_ and *b*_*k*_, and setting *γ*_*k*_ = 1 in the final expression. Derivation of the cfGMM parameter estimator can be found in the Supplementary Material Section 1. Note that the closed-form estimators are not identical to the MLE of the Gamma distribution, but they approach the MLE in large samples seen in mIF datasets (see Supplementary Material Section 3).

Sometimes, the local maxima of unsupervised clustering algorithms are not biologically reasonable because the marker positive and marker negative populations overlap. To address this issue, we add constraints on the mode of the components of the cfGMM in order to restrict the marker-positive cells to the right of the marker distribution. The constraints can be selected based on a percentile of the marker distribution or by visualizing the “elbow” of the expression distribution of the marker (see *GammaGateR pipeline*). To incorporate this biological information, we constrain the mode, *m*_*k*_, for component *k*, which is a function of parameters *a*_*k*_ and *b*_*k*_, so that it lies within the interval (*l*_*k*_, *u*_*k*_):

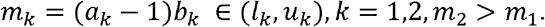

Because the the log-likelihood is strictly concave with respect to *a*_*k*_ and *b*_*k*_, the constrained maximum must lie on the boundary if the global maximum is outside the bounds, so finding the maximum reduces to finding the solution incorporating the boundary constraint. In this case, the expected log-likelihood of the component *k* is maximized on the line corresponding to the lower or upper constraint, e.g. (*a*_*k*_ − 1)*b*_*k*_ = *l*_*k*_. The parameters are estimated following the Expectation-Maximization algorithm [37] (See Supplementary Material Section 2).

#### GammaGateR pipeline

The analysis pipeline is illustrated for the CD4 marker channel (Figure 1). GammaGateR takes a single-cell image dataset, with each row corresponding to an individual cell, and each column as the normalized intensity of a marker channel for each cell (Figure 1.1). The first step is selecting biological constraints for model fitting by visualizing overlay histograms for each marker channel (Figure 1.2). The constraints are not manual thresholds, but represent boundaries for the mode of each component of the fitted distribution across all slides in the dataset. Because marker-positive cells often are a small proportion of all cells and have higher expression values, we limit the mode of the higher component to be no lower than the “elbow” of the overlay histograms (Figure 1.2, e.g. 0.45). While GammaGateR can be fit without constraints, the constraints provide more consistent model estimation across many slides. Examples of initial constraints for the three datasets presented here are given in Tables S3 & S4. Given the data and constraints, GammaGateR generates the parameter estimates of the Gamma mixture model including modes and proportion of each component, and the posterior and marginal probabilities of each cell being marker-positive.

**Figure 1:**
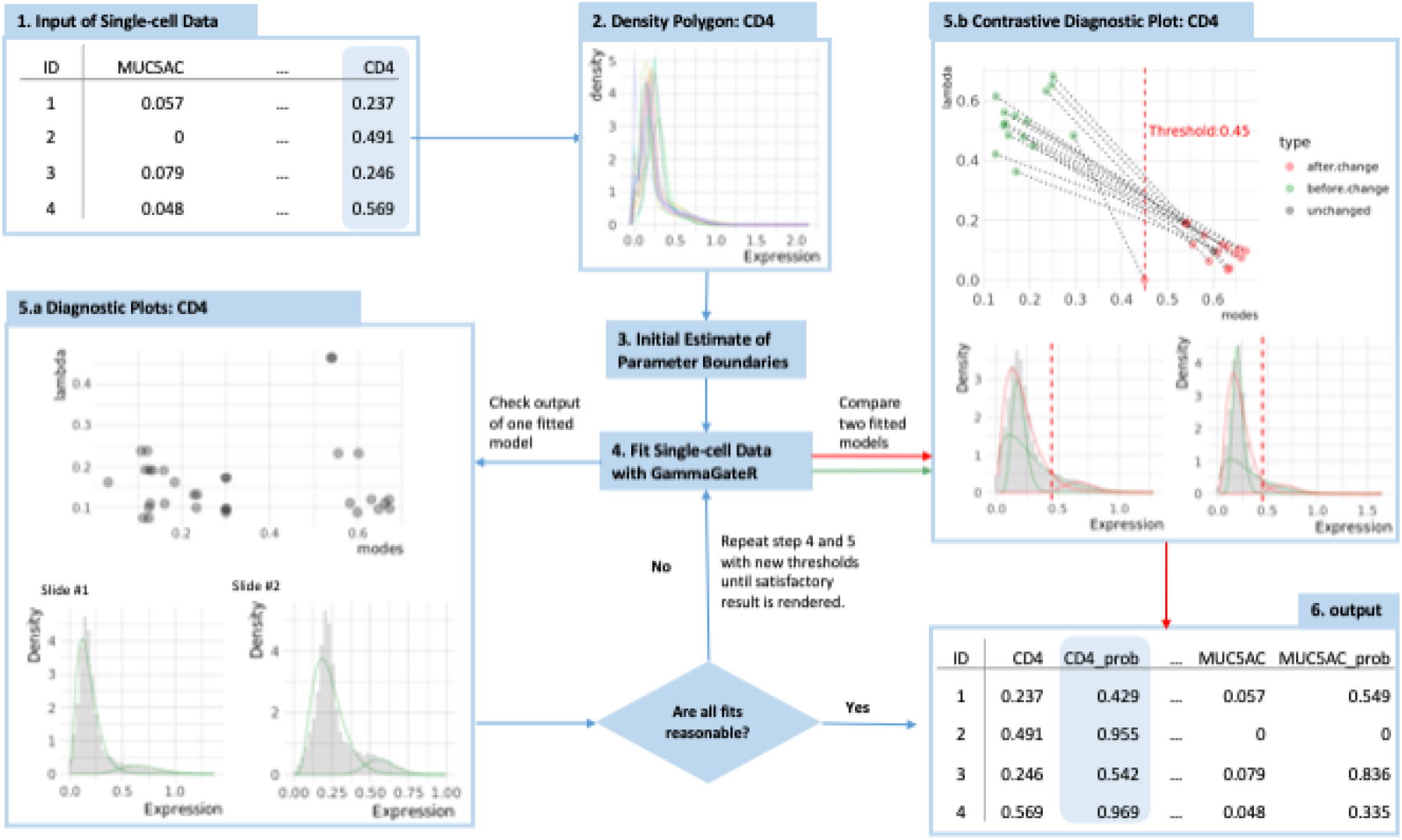
Overview of GammaGateR analysis pipeline for the CD4 marker channel. (1) GammaGateR takes segmented cell-level data as input. (2) Density polygons are used to visualize all slide level histograms and select constraints for model fit (3). (4) After model estimation, (5.a) diagnostic plots are used to evaluate the model fit. (5.b) New constraints can be selected and the refitted model can be compared to a previous model. (6) Expression probabilities can be extracted for downstream analysis from the model objects.

To ensure accurate model fitting, GammaGateR includes functionality for users to evaluate model fit and modify the fit when needed. Diagnostic plots for the fitted GammaGateR model object consist of a scatter plot of all the slides fitted model modes and lambda (marker-positive probability), and the fitted density curve over the cell expression histogram for each slide in the data set (Figure 1.5.a). The x-axis of the scatter plot is the mode for the marker-positive component, and the y-axis is the proportion of the corresponding component. The scatter plot is useful for identifying slides that are outliers with respect to where the mode of marker-positive cells lies or the estimated proportion of marker-positive cells. The histograms can be used for visually evaluating model fit for one slide. A good model fit shows an approximate fit of the smooth density line to the histogram with a marker-positive cell distribution sitting to the right (Figure 1.5.a). If there is poor model fit, users can compare fitted models between two different constraints to check how different boundaries affect fitted values (Figure 1.5.b). Figure 1.5.b compares the model fit for CD4 with no constraints (green) to the model fit with an initial constraint with a lower bound of 0.45 for the marker-positive component (red). The model without constraints places the marker-positive distribution directly over the marker-negative distribution. Users can adjust the parameter boundaries and fit again until satisfactory fittings are rendered. Finally, the output of the fitted models is easily accessible in the GammaGateR model object (Figure 1.6). The vignette in the GammaGateR R package provides a guide to fitting the GammaGateR model in the lung cancer dataset from the VectraPolaris dataset available on Bioconductor.

### 2.3 Model estimation

GammaGateR is fitted to all datasets following the procedure described in Figure 1. Among them, both the colon map data and the CRC atlas data go through an additional modification of the thresholds by increasing the thresholds after visually evaluating the model fit (Table S4). The ovarian cancer data shows good results in the first round of fitting. There are five slides that have one channel that did not converge in the colon map data and the CRC atlas data. These slides are discarded in the subsequent analyses. The markers, thresholds, adjusted thresholds, and number of slides that do not converge are all listed in Table S4.

We compare GammaGateR results to ASTIR, another unsupervised cell-level phenotyping software developed for mIF data [30]. ASTIR uses a combination of neural networks within a multivariate normal mixture model to obtain cell phenotypes from marker channel expression data and returns a vector of phenotype probabilities that sums to one for each cell. ASTIR takes the marker expression of each cell and an XML file as input to specify phenotypes based on prespecified marker combinations. The phenotypes input for ASTIR for the three datasets are given in Tables S1 and S2.

### 2.4 Methods and phenotyping evaluation

To compare the methods in determining cell phenotypes we assess the accuracy of each method relative to a “silver standard” manual phenotyping in the Colon MAP and CRC atlas datasets and evaluate the efficacy in predicting survival in the ovarian cancer data. We compare phenotyping results obtained using GammaGateR and ASTIR to “silver standard” manual phenotyping using the Adjusted Rand Index [38]. Adjusted rand index typically takes values between 0 and 1, where a larger value indicates a better alignment between two categorical variables. The silver standard is obtained by gating the raw images based on visual inspection and defining marker-positive cells as those that contain more than 50% marker-positive pixels within each cell. Semi-automated marker gating results are obtained using GammaGateR as described in Section 2.3, with monotonically adjusted posterior probability and marginal probability thresholded at 0.5 to define marker positive cells. The same phenotype definitions used for ASTIR are used to define phenotypes from marker positive labels using GammaGateR and manual marker gating as well [5,7]. Each cell belongs to a given phenotype if it is marker positive for combinations of markers for that phenotype (Tables S1, S2). ASTIR phenotypes are determined by selecting the cell type with the maximum probability for each cell. All methods use the same combinations of markers to define phenotypes (Table S1, S2).

Because the Ovarian Cancer dataset does not include manual cell phenotypes, we instead compare the prediction accuracy of survival time data for each method across all patients in the study to determine if one method has greater biological sensitivity than the other. The original study shows that survival of ovarian cancer patients is significantly correlated with B-cell and CD4 T-cell, as well as spatial interaction between CD4 T-cell and macrophage. Therefore, we evaluate the methods using this dataset by fitting a survival model with age, cancer stage, B-cell proportions, CD4 T-cell proportions, and the spatial interaction between Macrophage and CD4 T-cells estimated by Ripley’s H with r=50 [39]. Ripley’s H is a geospatial index that describes spatial attraction/repulsion. We fit a model for each method, where the cell phenotypes are determined using the given method, and compare all models, as well as a base model that includes only age and cancer stage. We use a random forest survival model to be sensitive to complex nonlinear relationships [40,41]. To estimate variability in the out-of-bag performance error, we compare the methods across 100 bootstrap samples. Performance error quantification is based on C-index [42], where a low performance error means that the model is a good fit, and a performance error of 0.5 is a random chance.

### 2.6 Spatial analysis in Colon MAP

We use GammaGateR to perform analyses of spatial features of the tumor microenvironment in precancerous colon polyps using the Colon MAP dataset [5]. Specifically, we summarize immune cell population proportions (CD3+) and MUC5AC expression in the manually annotated tumor region of each sample, as well as quantify the spatial interaction between MUC5AC+ cells and CD68+ cells. To estimate cell population proportions, instead of dichotomizing cell populations, we average the posterior marker positive probabilities across all cells in the tumor region. For example, the proportion of CD3+ cells in the tumor epithelium is quantified as

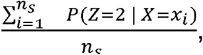

where *x*_*i*_ is the CD3 marker value for cell *i*, in the tumor epithelial region, *n*_*s*_ is the total number of cells segmented in the tumor epithelial slide *s*, and *P*(*Z* = 2 | *X* = *x*_*i*_) is as described in Equation (2). To study differences in cell proportions between tumor types we fit a linear regression model on the logit transformed probabilities with regions clustered within samples and weights proportional to the number of cells in each region. We report hypothesis tests using robust standard errors and a robust effect size index (RESI) with 95% confidence intervals to quantify the effect of tumor types on marker channels [43,44].

To quantify spatial clustering, we threshold the posterior probabilities and use Ripley’s H with r ∈ (0,1000) to quantify spatial attraction or repulsion of MUC5AC+ and CD68+ cells within each region of each tissue sample [39]. After estimating Ripley’s H, we average the index across all radii and test the difference between groups using a Welch t-test.

## 3. Results

### 3.1 Performance Evaluation

We compare the ARI of each method relative to a silver standard manual phenotype in the Colon Map and CRC atlas datasets. For both datasets and all cell types in these datasets, the posterior probability by GammaGateR yields higher Median ARI (Figure 2a & b). This means that the posterior probability has consistently greater similarity to the silver standard than marginal probability and ASTIR (Figure 2a & b). However, for some cell types (e.g. Macrophage, B-cells, Myeloid), all methods have low performance. This is an indication of systematic difference in how the algorithms identify positive cells relative to the manual labels.

**Figure 2:**
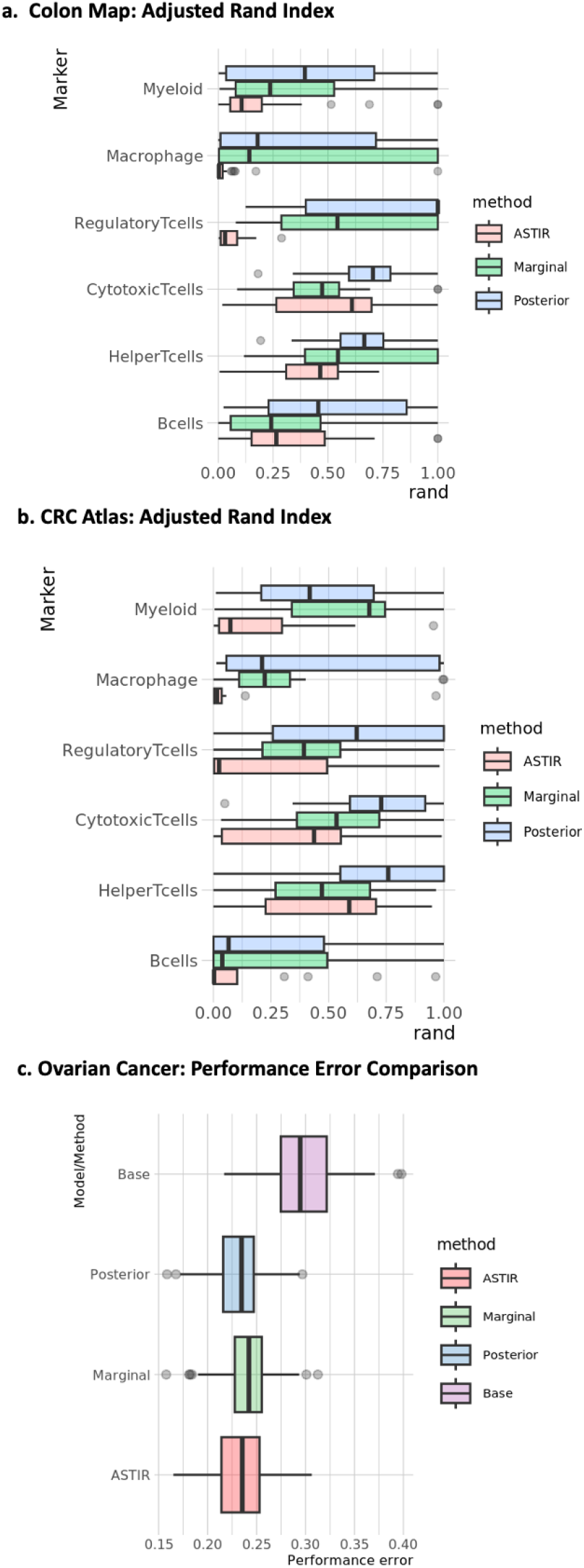
Performance evaluation for GammaGateR on the three datasets. “Posterior” and “marginal” refer to the posterior and marginal probabilities from GammaGateR, respectively. Cell phenotyping performance comparing GammaGateR to ASTIR in the (a) Colon MAP and (b) CRC Spatial atlas. (c) Survival prediction performance error in the ovarian cancer dataset, measured by 1-C-index. “Base” indicates the survival model including only age and cancer stage variables.

To evaluate the methods in the Ovarian cancer data in the absence of manual phenotypes, we use proportions and spatial characteristics of the phenotypes from each method as predictors in a random forest survival model. For all methods, incorporating the cell-level information reduces out-of-bag error performance by approximately 0.075, over the base model that includes only age and cancer stage. This indicates that spatial cell phenotype covariates are useful in predicting survival outcomes, consistent with the original findings [33]. The posterior probability slightly outperforms other methods, having the lowest prediction error in 46% of the bootstrap samples, compared to 36% with ASTIR and 18% with the marginal probability (Figure 2c).

### 3.2 Cell Spatial Interaction in Colon MAP

We use marker gating results from GammaGateR to compare spatial features across adenoma and SSL samples. We hypothesize that SSLs, which are precursors to CRCs with high microsatellite-instability that have better prognosis, have greater immune cell infiltration in the epithelial tissue (greater proportion of CD3+ cells), greater expression of MUC5AC in the tumor region, and that cells expressing MUC5AC spatially attract Macrophages (CD68+ cells) in SSLs, but not in adenoma samples [5].

From the results, we observe that SSLs are associated with enriched immune cell populations (CD3+ cell proportions) in the epithelial region of the tumor tissue (Figure 3a; T=2.22, df=35, p=0.0332, RESI=0.15, 95% CI for RESI=(0, 0.52)) and have higher, but a nonsignificant proportion of MUC5AC+ cells in the tumor region (Figure 3b; T=1.83, df=13, p=0.0902, RESI=0.20, 95% CI for RESI=(0, 0.97)). Ripley’s H shows a strong difference between AD and SSL slides (Figure 3c; T= -4.67, df = 8.56, p = 0.001, RESI=1.01 95% CI for RESI=(0.60, 1.87)). The values of Ripley’s H indicate spatial attraction between MUC5AC+ and CD68+ cells in SSLs, but spatial repulsion between these two in adenomas. These results are reflected visually in representative samples of the GammaGateR phenotypes with the raw imaging data (Figure 3c). The concordance between quantified measurements generated by GammaGateR and qualitative observation conclusions shows that GammaGateR can help solidify observed spatial pattern.

**Figure 3:**
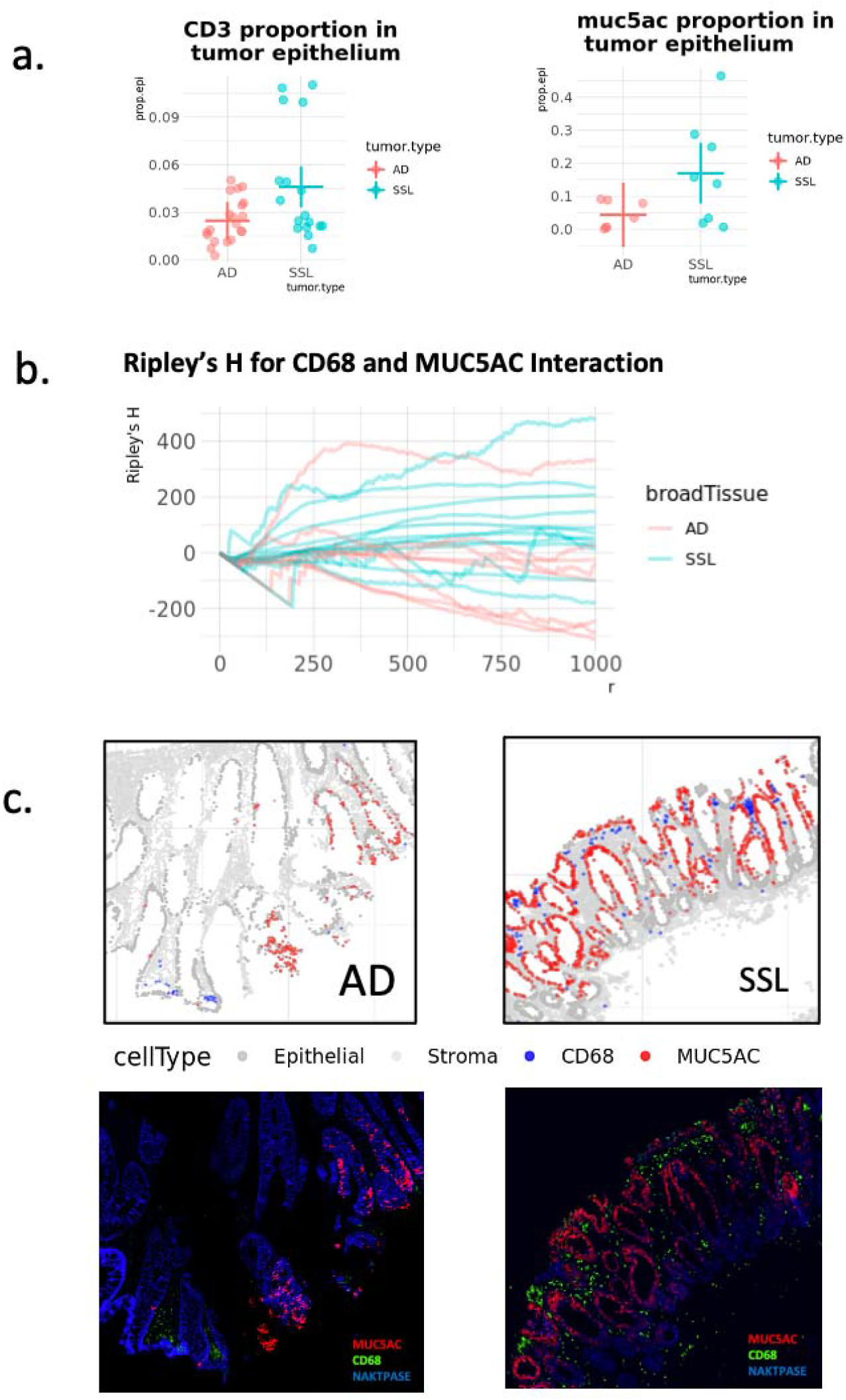
Spatial analysis results using GammaGateR in the Colon MAP data. “AD” and “SSL” represent adenoma and sessile serrated lesions, respectively. a) Comparison of CD3+ and MUC5AC+ cell proportions between tumor types in epithelial regions of the tumor mask. Each point is the mean marker positive probabilities for one slide. The horizontal lines are means and vertical lines are robust 95% confidence intervals. b) Ripley’s H curves for spatial interaction between MUC5AC+ and CD68+ cells for each slide. c) Examples of MUC5AC+ and CD68+ cells identified with GammaGateR in the two tumor types, with corresponding raw image intensities in the multiplex images (MUC5AC, CD68, and NAKATPASE).

## 4. Summary and Discussion

We introduced GammaGateR, a semi-automated maker gating tool. Driven by a novel cfGMM estimation framework, GammaGateR generates reproducible and evaluable marker gating for mIF data. In addition, cfGMM enables computationally feasible model estimation for large-scale datasets like mIF. The marker gating output of GammaGateR can be used to define phenotypes and as input to downstream analysis. GammaGateR implements interactive visualization to quality check the clustering results (see vignette in Supplementary Material) and allows users to modify the constraints to improve model results when needed. Consequently, GammaGateR provides more consistent results with silver standard labels than ASTIR, the existing state-of-the-art method for automated phenotyping of cell-level mIF data. In the examples shown in this paper, GammaGateR phenotypes had slightly improved ovarian cancer survival prediction accuracy compared to ASTIR. This paper also compares the posterior and the marginal probabilities returned by GammaGateR. The marginal probabilities only use the marker positive cell distribution to determine cell phenotypes, whereas the posterior probabilities take into account the distribution of the marker negative cells. Using posterior probabilities was almost always better indicating the importance of accounting for the full distribution of the marker intensities when identifying marker-positive cells.

Despite being an effective tool with a high level of implementation, GammaGateR could still use a few improvements to broaden its usage. First, the accuracy of GammaGateR might further improve with pixel-level information. GammaGateR is applied to segmented cell-level data and does not rely on patterns of pixel intensities within and around each cell. While this makes GammaGateR easy to apply in many datasets, it obviously does not leverage the information available at the pixel level around each cell, which may be useful for identifying doublets and improving cell phenotype identification as in recent work [45,46]. Second, GammaGateR focuses on obtaining a highly accurate fit to the marginal distribution of each marker channel for each slide and is applied separately to each slide and channel making it highly parallelizable. One disadvantage of this approach is that it does not explicitly account for the joint distributions of markers, which could be used for obtaining more accurate phenotypes. Instead, multiple markers can be incorporated by combining the posterior probabilities to define phenotypes as was performed above in the Colon MAP and CRC atlas datasets. Future work could model the multivariate distributions. However, it should be noted that this approach risks phenotypes being driven, in part, by channels that might not be relevant for a given cell type. Third, more thorough evaluation of batch effect could render more reasonable marker-gating output. We applied normalization prior to running GammaGateR to reduce the influence of batch effects. Although GammaGateR does not explicitly model and remove batch effects, because the model is fit separately for each slide, it may help to mitigate batch effects. When using the posterior probabilities in downstream analysis an implicit assumption is that differences in the shape and location of the marker positive component do not represent biological differences. Finally, in our applications, five models had too few cells to estimate or did not converge and were excluded from the analysis. GammaGateR parameter estimates for the failed models could be imputed by combining estimates across slides, which might also help with the estimation and removal of batch effects.

GammaGateR is the first semi-automated marker gating method developed specifically for mIF data and is useful to define quality-controlled marker-positive cells by flexibly modeling marker distributions across cells and channels. GammaGateR has demonstrated consistency with manual labels and sensitivity to biological information which makes it another useful method for the multiplexed imaging scientist.

## DATA AVAILABILITY STATEMENT

The R package GammaGateR can be installed through Github (https://github.com/JiangmeiRubyXiong/GammaGateR). The R package cfGMM can be installed through Github (https://github.com/JiangmeiRubyXiong/cfGMM).

## ACKNOWLEDGMENTS

We would like to thank Dr.Diane Saunders, Conrad Reihsmann, and Alexander Hopkirk from Vanderbilt Diabetes Research Center for their generous help with data analysis.

## Table Captions

Table S1. Marker-to-phenotype correspondence in colon precancer and CRC atlas datasets.

Table S2. Marker-to-phenotype correspondence in ovarian cancer dataset.

Table S3. Examples of initial constraints for the ovarian cancer data.

Table S4. Constraints, adjusted constraints, and convergence for the colon map data and CRC atlas data. GammaGateR was only fit on the subset of markers used in the paper for the CRC atlas.

