## Supplementary Material for "GammaGateR: semi-automated marker gating for single-cell multiplexed imaging"

August 2023

#### 1 Derivation of cfGMM estimator

We assume the data is a random sample $x_1, \ldots, x_n$ from a $K$ component generalized gamma mixture distribution. The density function of $X$ is

$$P(X = x) = \sum_{k=1}^{K} \lambda_k f(x; a_k, b_k, \gamma_k).$$

and the log-likelihood of the dataset is

$$\ell(\mathbf{x}|\mathbf{a}, \mathbf{b}, \boldsymbol{\lambda}) = \sum_{i=1}^{n} \log \left\{ \sum_{k=1}^{K} \lambda_k f(x_i|a_k, b_k) \right\} \tag{1}$$

For each generalized gamma component $k$, $\lambda_k \in [0, 1]$ are the mixture parameters, $\sum_k \lambda_k = 1$; $f$ denotes the generalized gamma density function; $a_k, b_k, \gamma_k$ are the parameters for the generalized gamma.

Here, we use the expectation maximization (EM) algorithm [Dempster et al., 1977] for parameter estimation. EM algorithm is a standard approach for parameter estimation in mixture models. It introduces the latent multinomial variable $Z_i = (Z_{i1}, \ldots, Z_{iK})$ into the model and maximizes the expected value of the complete data likelihood[Dempster et al., 1977]. The expectation of the complete data likelihood to be maximized for the generalized gamma distribution is

$$\mathbb{E}_Z \ell(x \mid Z) = \sum_{i=1}^{n} \sum_{k=1}^{K} z_{ik} \log f(x_i; a_k, b_k, \gamma_k),$$

where

$$z_{ik} = \mathbb{P}(Z_{ik} = 1 \mid x_i; \boldsymbol{a}, \boldsymbol{b}, \boldsymbol{\gamma}) = \frac{f(x_i|a_k, b_k, \gamma_k)}{\sum_{j=1}^{K} f(x_i|a_j, b_j, \gamma_k)}, \tag{2}$$

$\mathbf{a} = (a_1, a_2, \ldots, a_K)$, and $\mathbf{b}$, $\boldsymbol{\lambda}$ are similarly defined vectors.

From here, the maximization of the expectation is now analogous to the maximization of generalized gamma distribution for each component of the mixture model.

The expectation of log-likelihood is

$$\mathbb{E}_{z|x}[\log(L(\mathbf{x}|\mathbf{z}))] = \sum_{i=1}^{n} \sum_{k=1}^{K} z_{ik} \log f_k(x_i) \tag{3}$$

and

$$f_k(x) = G(a_k, b_k, \gamma_k) = \frac{\lambda_k x^{a_k\gamma_k - 1} exp\{(-x/b_k)^{\gamma_k}\}}{b_k^{a_k\gamma_k}\Gamma(a_k)} \tag{4}$$

where $\gamma_k = 1$.

By (3), (4) The expected joint log-likelihood is

$$\mathbb{E}_{z|x}[\log(L(\mathbf{x}|\mathbf{z}))] \quad = \quad \sum_{k=1}^{K}\sum_{i=1}^{n} z_{ik}\left(\log\gamma_k - a_k\gamma_k\log b_k - log\Gamma(a_k) + (a_k\gamma_k - 1)\log X_i - (\frac{X_i}{b_k})^{\gamma_k}\right) \tag{5}$$

The estimators of each of the $K$ terms of the expected joint log-likelihood are derived as follows: first take derivative of the expression from (5)

$$\sum_{i=1}^{n} z_{ik}\left(\log\gamma_k - a_k\gamma_k\log b_k - \log\Gamma(a_k) + (a_k\gamma_k - 1)\log X_i - \left(\frac{X_i}{b_k}\right)^{\gamma_k}\right)$$

with respect to $a_k, b_k, \gamma_k$ separately:

$$\frac{\partial\mathbb{E}_{z|x}[\log(L(\mathbf{x}|\mathbf{z}))]}{\partial a_k} = \sum_{i=1}^{n} z_{ik}(-\psi(a_k) - \gamma_k\log b_k + \gamma_k X_i) = 0 \tag{6}$$

Note that $\psi(x) = \frac{d}{dx}\log\Gamma(x)$ is digamma function.

$$\frac{\partial\mathbb{E}_{z|x}[\log(L(\mathbf{x}|\mathbf{z}))]}{\partial b_k} = \sum_{i=1}^{n}(z_{ik})(-a_k\gamma_k/b_k + \gamma_k X_i^{\gamma_k} b_k^{-\gamma_k - 1}) = 0 \tag{7}$$

$$\frac{\partial\mathbb{E}_{z|x}[\log(L(\mathbf{x}|\mathbf{z}))]}{\partial \gamma_k} = \sum_{i=1}^{n} z_{ik}\left(\frac{1}{\gamma_k} - a_k\log b_k + a_k\log X_i - \left(\frac{X_i}{b_k}\right)^{\gamma_k}\log\frac{X_i}{b_k}\right) = 0 \tag{8}$$

Among which, (7) can be solved as

$$\hat{b}_k(a_k, \gamma_k) = \left(\frac{\sum_{i=1}^{n} z_{ik}X_i^{\gamma_k}}{a_k\sum_{i=1}^{n} z_{ik}}\right)^{1/\gamma_k} \tag{9}$$

Substitute (9) into (8):

$$\frac{\partial \mathbb{E}_{z|x}[\log(L(\mathbf{x}|\mathbf{z}))]}{\partial \gamma_k}$$

$$= \sum_{i=1}^{n} z_{ik}/\gamma_k + \sum_{i=1}^{n} a_k z_{ik} \log(\frac{X_i}{b_k}) - \sum_{i=1}^{n} z_{ik}(\frac{X_i}{b_k})^{\gamma_k} \log(\frac{X_i}{b_k})$$

$$= \sum_{i=1}^{n} z_{ik}/\gamma_k + \sum_{i=1}^{n} a_k z_{ik}(\log X_i - \log b_k) - b_k^{-\gamma_k} \sum_{i=1}^{n} z_{ik} X_i^{\gamma_k}(\log X_i - \log b_k)$$

$$= \sum_{i=1}^{n} z_{ik}/\gamma_k + \sum_{i=1}^{n} a_k z_{ik} \log X_i - \log b_k \sum_{i=1}^{n} a_k z_{ik} - b_k^{-\gamma_k} \sum_{i=1}^{n} z_{ik} X_i^{\gamma_k} \log X_i + b_k^{-\gamma_k} \log b_k \sum_{i=1}^{n} z_{ik} X_i^{\gamma_k}$$

$$= \sum_{i=1}^{n} z_{ik}/\gamma_k + \sum_{i=1}^{n} a_k z_{ik} \log X_i - \log b_k \sum_{i=1}^{n} a_k z_{ik} - b_k^{-\gamma_k} \sum_{i=1}^{n} z_{ik} X_i^{\gamma_k} \log X_i \frac{a_k \sum_{i=1}^{n} z_{ik}}{\sum_{i=1}^{n} z_{ik} X_i^{\gamma_k}} \log b_k \sum_{i=1}^{n} z_{ik} X_i^{\gamma_k}$$

$$= \sum_{i=1}^{n} z_{ik}/\gamma_k + \sum_{i=1}^{n} a_k z_{ik} \log X_i - \log b_k \sum_{i=1}^{n} a_k z_{ik} - b_k^{-\gamma_k} \sum_{i=1}^{n} z_{ik} X_i^{\gamma_k} \log X_i + a_k \sum_{i=1}^{n} z_{ik} \log b_k$$

$$= \sum_{i=1}^{n} z_{ik}/\gamma_k + a_k \sum_{i=1}^{n} z_{ik} \log X_i - \frac{a_k \sum_{i=1}^{n} z_{ik}}{\sum_{i=1}^{n} z_{ik} X_i^{\gamma_k}} \sum_{i=1}^{n} z_{ik} X_i^{\gamma_k} \log X_i$$

$$= \sum_{i=1}^{n} z_{ik}/\gamma_k + a_k \left( \sum_{i=1}^{n} z_{ik} \log X_i - \frac{\sum_{i=1}^{n} z_{ik}}{\sum_{i=1}^{n} z_{ik} X_i^{\gamma_k}} \sum_{i=1}^{n} z_{ik} X_i^{\gamma_k} \log X_i \right) = 0$$

Solving this, we have

$$\hat{a}_k(\gamma_k) = \frac{\sum_{i=1}^{n} \frac{z_{ik}}{\gamma_k}}{\frac{\sum_{i=1}^{n} z_{ik}}{\sum_{i=1}^{n} z_{ik} X_i^{\gamma_k}} \sum_{i=1}^{n} z_{ik} X_i^{\gamma_k} \log X_i - \sum_{i=1}^{n} z_{ik} \log X_i} \tag{10}$$

Plug $\gamma_k = 1$ in (10), we now have

$$\hat{a}_k(\gamma_k = 1) = \frac{\sum_{i=1}^{n} z_{ik}}{\frac{\sum_{i=1}^{n} z_{ik}}{\sum_{i=1}^{n} z_{ik} X_i} \sum_{i=1}^{n} z_{ik} X_i \log X_i - \sum_{i=1}^{n} z_{ik} \log X_i}$$

$$= \left( \frac{\sum_{i=1}^{n} z_{ik} X_i \log X_i}{\sum_{i=1}^{n} z_{ik} X_i} - \frac{\sum_{i=1}^{n} z_{ik} \log X_i}{\sum_{i=1}^{n} z_{ik}} \right)^{-1}$$

$$= \frac{\sum_{i=1}^{n} z_{ik} \sum_{i=1}^{n} z_{ik} X_i}{\sum_{i=1}^{n} z_{ik} \sum_{i=1}^{n} z_{ik} X_i \log X_i - \sum_{i=1}^{n} z_{ik} \log X_i \sum_{i=1}^{n} z_{ik} X_i} \tag{11}$$

$$\hat{b}_k(\hat{a}_k, \gamma_k = 1) = \frac{\sum_{i=1}^{n} z_{ik} X_i}{\hat{a}_k \sum_{i=1}^{n} z_{ik}}$$

$$= \frac{\sum_{i=1}^{n} z_{ik} \sum_{i=1}^{n} z_{ik} X_i \log X_i - \sum_{i=1}^{n} z_{ik} \log X_i \sum_{i=1}^{n} z_{ik} X_i}{\left( \sum_{i=1}^{n} z_{ik} \right)^2} \tag{12}$$

In addition, $\hat{\lambda}_k$ can simply be estimated as

$$\hat{\lambda}_k = \frac{\sum_{i=1}^{n} z_{ik}}{n} \tag{13}$$

It is worth noting that we are not maximizing the exact Gamma distribution, therefore the algorithm we devise here is an EM-type algorithm. Procedure 2.1 shows how it is carried out.

### 2 Fitting with constraints

In many applications, we have prior information that the mode of components are within a certain area. For example, in mIF imaging data we know that, for the CD8 marker, there is a small proportion of CD8+ cells whose mode should be to the right of the unexpressed cells. To incorporate this information, we constrain the mode, $m_k$, which is a function of parameters $a_k, b_k$, to be in an interval

$$m_k = (a_k - 1)\frac{1}{b_k} \in (l_k, u_k), k = 1, 2, 3, ..., K \tag{14}$$

This constraint is equivalent to putting a flat bounded prior on the mode of the components. Because the the log-likelihood is strictly concave with respect to $a_k$ and $b_k$, the constrained maximum must lie on the boundary, if the global maximum is outside the bounds [Boos and Stefanski, 2013]. In this case, the expected log-likelihood of the component $k$ is maximized at $m_k$

$$(a_k - 1)\frac{1}{b_k} = m_k \Rightarrow a_k = m_k \frac{1}{b_k} + 1 \tag{15}$$

Figure 1 is a graphical depiction of the boundaries and likelihood, where the likelihood is generated from one of 2-component datasets from the simulation section. To find the constrained MLE of $b_k$, we can plug equation (15) in the expected log-likelihood with $\gamma_k = 1$, take derivative with respect to $b_k$, and set it equal to zero

$$\frac{\partial \mathbb{E}_{z|x}[\log(L(\mathbf{x}|\mathbf{z}))]}{b_k} = \sum_{i=1}^{n} \frac{z_{ik}}{-b_k^2}\Big(m_k + b_k - m_k \log(b_k) - m_k\psi(m_k/b_k + 1) + m_k \log X_i - X_i.\Big) = 0 \tag{16}$$

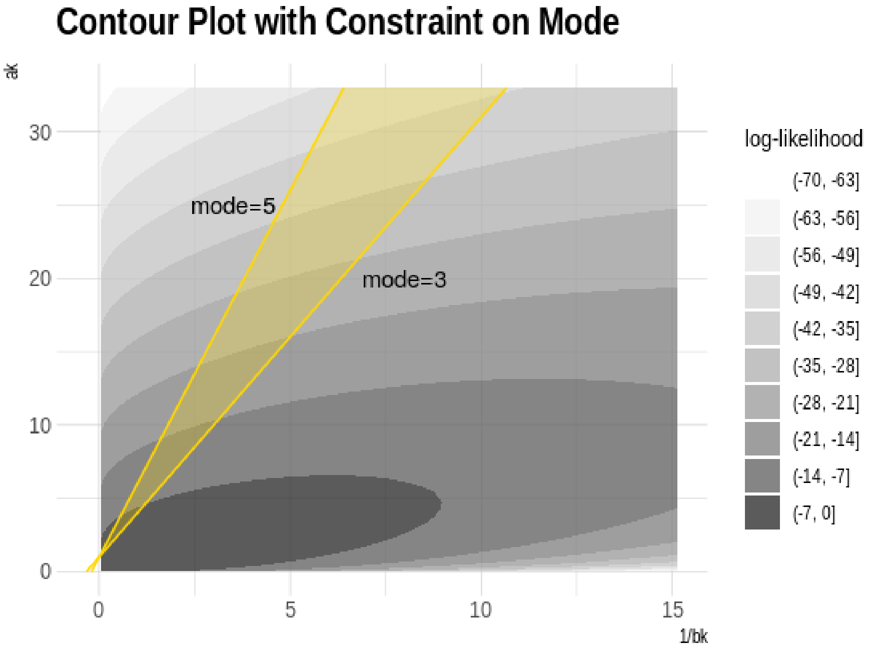


Figure 1: A contour plot of log-likelihood with mode constraints. The log-likelihood is created using one of the two-component simulation data sets. The straight lines are the $m_k = 3, m_k = 5$ boundary, separately and their expression follows equation (15). The range allowed for mode is the shaded area between the line of mode bounds. Due to concavity of the likelihood function the constrained maximum occurs on the boundary.

Unfortunately, this equation does not have a closed-form solution due to the digamma function, $\psi(.)$. Therefore, we use Newton-Raphson to solve for the optimal $b_k$. This approach gives Procedure 2.2.

#### 2.1 EM-type Algorithm Procedure

**Procedure 2.1 (cfGMM EM-type algorithm)** *Let $K$ denote the number of components, $M$ denote the maximum number of iterations, and $\epsilon > 0$ be the convergence criterion.*

1. *Initialize* $C = \infty$*,* $t = 1$*. Randomly generate* $\boldsymbol{\lambda}^{(0)}$ *so that* $\sum_k \lambda_k = 1$*. Initialize parameter vectors of* $\boldsymbol{a}^{(0)}, \boldsymbol{b}^{(0)} \in \mathbb{R}^K$ *using method of moments (MOM) estimator of the data partitioned according to* $\boldsymbol{\lambda}^{(0)}$*.*

2. *While* $t < M$ *and* $C > \epsilon$*, compute the expected values* $z_{ik}^{(t)}$ *using formula* (2)*, with* $\boldsymbol{a} = \boldsymbol{a}^{(t-1)}$ *and* $\boldsymbol{b} = \boldsymbol{b}^{(t-1)}$*. Then compute* $\boldsymbol{a}^{(t)}$*,* $\boldsymbol{b}^{(t)}$ *and* $\boldsymbol{\lambda}^{(t)}$ *that maximize the expected log-likelihood using formulas* (11)*,* (12) *and* (13) *with* $z_{ik} = z_{ik}^{(t)}$*. Use formula* (1) *to set* $C = \left|\ell(\mathbf{x}|\mathbf{a}^{(t)}, \mathbf{b}^{(t)}, \boldsymbol{\lambda}^{(t)}) - \ell(\mathbf{x}|\mathbf{a}^{(t-1)}, \mathbf{b}^{(t-1)}, \boldsymbol{\lambda}^{(t-1)})\right|/n$*,* $t = t + 1$*.*

3. *Set* $\hat{\boldsymbol{a}} = \boldsymbol{a}^{(t)}, \hat{\boldsymbol{b}} = \boldsymbol{b}^{(t)}$*, and* $\hat{\boldsymbol{\lambda}} = \boldsymbol{\lambda}^{(t)}$*.*

**Procedure 2.2 (constrained cfGMM EM-type algorithm)** *Let* $K$ *denote the number of components,* $M$ *denote the maximum number of interations, and* $\epsilon > 0$ *be the convergence criterion.*

1. *Initialize* $C = \infty$*,* $t = 1$*. Randomly generate* $\boldsymbol{\lambda}^{(0)}$ *so that* $\sum_k \lambda_k = 1$*. Initialize parameter vectors of* $\boldsymbol{a}^{(0)}, \boldsymbol{b}^{(0)} \in \mathbb{R}^K$ *using method of moments (MOM) estimator of the data partitioned according to* $\boldsymbol{\lambda}^{(0)}$*.*

2. *While* $t < M$ *and* $C > \epsilon$
   (a) *compute* $z_{ik}^{(t)}$ *using formula* (2)*, with* $\boldsymbol{a} = \boldsymbol{a}^{(t-1)}$ *and* $\boldsymbol{b} = \boldsymbol{b}^{(t-1)}$*.*
   (b) $\boldsymbol{a}^{(t)}$*,* $\boldsymbol{b}^{(t)}$ *and* $\boldsymbol{\lambda}^{(t)}$ *using formulas* (11)*,* (12) *and* (13) *with* $z_{ik} = z_{ik}^{(t)}$*. Check if equation* (14) *holds true for all pairs of* $a_k^{(t)}, b_k^{(t)}$*. If not, set* $m_k$ *to its closest boundary, and plug it in equation* (16) *to solve for* $b_k$*, and set* $a_k$ *by equation* (15)
   (c) *Using formula* (1)*, set* $C = \left|\ell(\mathbf{x}|\mathbf{a}^{(t)}, \mathbf{b}^{(t)}, \boldsymbol{\lambda}^{(t)}) - \ell(\mathbf{x}|\mathbf{a}^{(t-1)}, \mathbf{b}^{(t-1)}, \boldsymbol{\lambda}^{(t-1)})\right|/n$ *and* $t = t + 1$*.*

3. *Set* $\hat{\boldsymbol{a}} = \boldsymbol{a}^{(t)}, \hat{\boldsymbol{b}} = \boldsymbol{b}^{(t)}$*, and* $\hat{\boldsymbol{\lambda}} = \boldsymbol{\lambda}^{(t)}$

### 3 Simulation

#### 3.1 Simulation setup

To compare the bias and compute time of the closed-form GMM to maximum likelihood GMM implementation, we run the cfGMM, the constrained cfGMM, and the GMM to evaluate bias and variance in a sample size of 10,000 across 1,000 simulations. We simulate a two-component mixture model with parameters $\boldsymbol{\lambda} = (0.3, 0.7), \boldsymbol{a} = (0.5, 8), \boldsymbol{b} = (0.5, 1/3)$. For the constrained estimator, we restrict the mode of each component to be in the range $(-\infty, 0)$ and $(0, 5)$ for marker negative and marker positive components, respectively, which include the true mode for each component, 0 (no mode) and 7/3.

#### 3.2 Simulation analysis

We use simulations to assess the bias and computing time of the closed-form model estimation in comparison to the MLE. Both closed-form estimation procedures have substantially faster computation time than the MLE (Figure 2a) while maintaining similarly low bias (Figure 2b). The sample size used in the simulation is roughly similar to that of the cell-level mIF image dataset, which further proves that cfGMM brings computation efficiency to our target application. The closed-form GMM, therefore, enables computationally feasible, precise, and flexible model estimation when applied to a large number of channels and slides using GammaGateR. It is also worth noting that the constrained cfGMM converges slightly faster than without constraints. This implies that when using cfGMM, computational cost can be reduced with proper knowledge of biological priors.

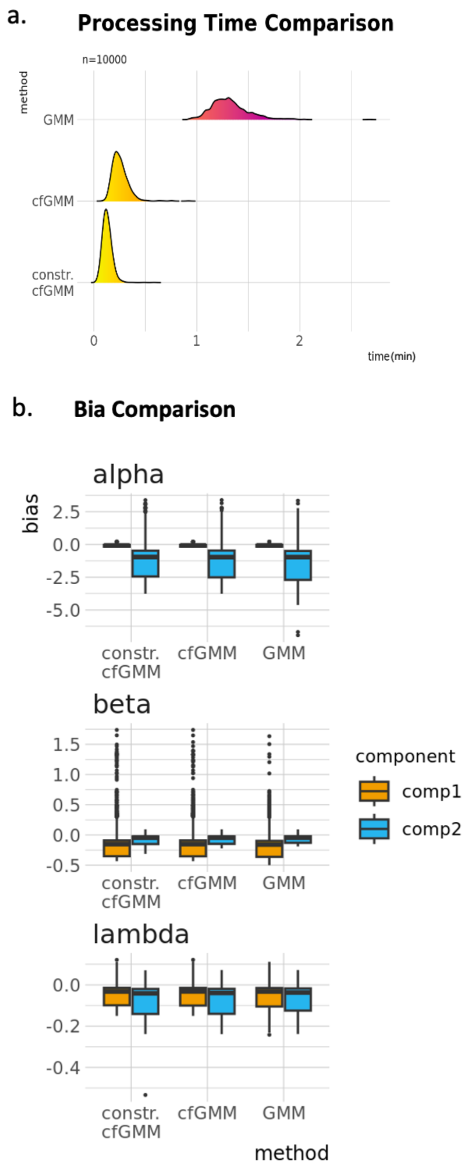


Figure 2: Simulation results for cfGMM performance evaluation. a) Run time comparison (in minutes) for three methods: GMM, cfGMM and cfGMM with constraints. b) Estimated bias across 1,000 simulations in a sample size of 10,000.
