## Supplementary material for "GammaGateR: semi-automated marker gating for single-cell multiplexed imaging": Table S

Supplementary Material – Tables

Table S1. Marker-to-phenotype correspondence in colon precancer and CRC atlas datasets.

| Cell Types | Markers |
| --- | --- |
| B cells | CD20+ |
| Helper T cells | CD3D+ CD4+ FOXP3- |
| Cytotoxic T cells | CD3D+ CD8+ |
| Regulatory T cells | CD3D+ CD4+ FOXP3+ |
| Macrophage | CD11B+  CD68+ LYSOZYME+ |
| Myeloid | CD11B+  (CD68-/ LYSOZYME-) |

Table S2. Marker-to-phenotype correspondence in ovarian cancer dataset.

| Cell Types | Markers |
| --- | --- |
| B cells | CD19+ |
| Macrophage | CD68+ |
| CD8 T cells | CD3+ CD8+ |
| CD4 T cells | CD3+ CD8- |
| Tumor | CK |

Table S3. Examples of initial constraints for the ovarian cancer data.

| Marker | Boundary - Component 1 | Boundary - Component 2 |
| --- | --- | --- |
| ck | (0,0.1) | (0.1,1) |
| ki67 | (0,0.2) | (0.2,1) |
| cd8 | (0,0.5) | (0.5,1) |
| ier3 | (0,0.5) | (0.5,1) |
| pstat2 | (0,0.5) | (0.5,1) |
| cd3 | (0,0.5) | (0.5,1) |
| cd68 | (0,0.5) | (0.5,1) |
| cd19 | (0,0.5) | (0.5,1) |
| dapi | (0,0.5) | (0.5,1) |
| autofluorescence | (0,0.5) | (0.5,1) |

Table S4. Constraints, adjusted constraints, and convergence for the colon map data and CRC atlas data. GammaGateR was only fit on the subset of markers used in the paper for the CRC atlas.

| Marker | Colon Map Data | | | CRC Atlas Data | | |
| --- | --- | --- | --- | --- | --- | --- |
|  | Constraints  (Comp 1)  (Comp 2) | Adjusted Constraints | N Not converged | Constraints  (Comp 1)  (Comp 2) | Adjusted Constraints | N Not converged |
| BCATENIN | (-Inf, 0.25)  (0.25, Inf) |  | 0 |  |  |  |
| CD11B | (-Inf, 0.25)  (0.25, Inf) |  | 0 | (-Inf, 0.3) (0.25, Inf) | (-Inf, 0.3) (0.5, 1.05) | 0 |
| CD20 | (-Inf, 0.35)  (0.35, Inf) | (-Inf, 0.5)  (0.5, 0.8) | 0 | (-Inf, 0.4)  (0.4, Inf) | (-Inf, 0.54)  (0.54, 0.75) | 0 |
| CD3D | (-Inf, 0.3)  (0.3, Inf) | (-Inf, 0.5)  (0.5, Inf) | 1 | (-Inf, 0.3) (0.25, Inf) | (-Inf, 0.5) (1.5, Inf) | 0 |
| CD4 | (-Inf, 0.3)  (0.3, Inf) | (-Inf,.5)  (0 .5, Inf) | 1 | (-Inf, 0.4)  (0.4, Inf) | (-Inf, 0.4)  (0.4, 0.75) | 0 |
| CD45 | (-Inf, 0.25)  (0.25, Inf) | (-Inf, 0.5)  (0.5, Inf) | 2 |  | (-Inf, 0.5)  (0.5, Inf) | 0 |
| CD68 | (-Inf, 0.4)  (0.4, Inf) | (-Inf,.5)  (0.5, Inf) | 0 | (-Inf, 0.5)  (0 .5, Inf) |  | 0 |
| CD8 | (-Inf, 0.4)  (0.4, Inf) | (-Inf,.5)  (0.5, Inf) | 0 | (-Inf, 0.4) (0.5, Inf) | (-Inf, 0.4) (0.6, 1.1) | 1 |
| CGA | (-Inf, 0.4)  (0.4, Inf) | (-Inf,.4)  (0.5, .8) | 1 |  |  |  |
| COLLAGEN | (-Inf, 0.2)  (0.2, Inf) | (-Inf, 0.4)  (0.4, Inf) | 0 |  |  |  |
| DAPI |  |  |  | (-Inf, 0.4)  (0.4, Inf) |  | 0 |
| ERBB2 | (-Inf, 0.4)  (0.4, Inf) | (-Inf,.35)  (0.35, Inf) | 0 |  |  |  |
| FOXP3 | (-Inf, 0.05)  (0.05, Inf) | (-Inf, 0.5)  (0.5, Inf) | 0 |  | (-Inf, 0.3) (0.45, 1.5) | 0 |
| HLAA | (-Inf, 0.5)  (0.5, Inf) |  | 0 |  |  |  |
| LYSOZYME | (-Inf, 0.3)  (0.3, Inf) | (-Inf,.4)  (0.4, Inf) | 0 | (-Inf, 0.3)  (0.3, Inf) |  | 0 |
| MUC2 | (-Inf, 0.4)  (0.4, Inf) | (-Inf, 0.5)  (0.5, Inf) | 0 |  |  |  |
| NAKATPASE | (-Inf, 0.25)  (0.25, Inf) | (-Inf, 0.35)  (0.35, Inf) | 0 |  |  |  |
| OLFM4 | (-Inf, 0.4)  (0.4, Inf) | (-Inf, 0.5)  (0.5, Inf) | 0 |  |  |  |
| PANCK | (-Inf, 0.15)  (0.15, Inf) | (-Inf, 0.5)  (0.35, 0.6) | 0 |  |  | 0 |
| PCNA | (-Inf, 0.25)  (0.25, Inf) | (-Inf, 0.5)  (0.5, Inf) | 0 |  |  |  |
| PEGFR | (-Inf, 0.5)  (0.5, Inf) | (-Inf, 0.5)  (0.3, 0.7) | 0 |  |  |  |
| PSTAT3 | (-Inf, 0.5)  (0.5, Inf) | (-Inf, 0.5)  (0.5, 0.7) | 0 |  |  |  |
| SMA | (-Inf, 0.4)  (0.4, Inf) | (-Inf, 0.45)  (0.45, Inf) | 0 |  |  |  |
| SOX9 | (-Inf, 0.2)  (0.2, Inf) | -Inf, 0.3)  (0.3, Inf) | 0 |  |  |  |
| VIMENTIN | (-Inf, 0.1)  (0.1, Inf) | (-Inf, 0.3)  (0.3, Inf) | 0 |  |  |  |
| GACTIN | (-Inf, 0.5)  (0.5, Inf) | (-Inf, 0.5)  (0.1, Inf) | 0 |  |  |  |
| CDX2 | (-Inf, 0.1)  (0.1, Inf) | (-Inf, 0.5)  (0.5, Inf) | 0 |  |  |  |
| MUC5AC | (-Inf, 0.1)  (0.1, Inf) | (-Inf, 0.5)  (0.5, Inf) | 0 |  | (-Inf, 0.5)  (0.5, Inf) | 0 |
